# BandHiC: a memory-efficient and user-friendly Python package for organizing and analyzing Hi-C matrices down to sub-kilobase resolution

**DOI:** 10.1101/2025.10.16.682752

**Authors:** Weibing Wang, Junping Li, Yusen Ye, Lin Gao

## Abstract

Recent advances in high-resolution Hi-C and Micro-C technologies have enabled finer-scale characterization of 3D genome architecture, but they also introduce substantial computational challenges, as the size of dense contact matrices scales quadratically with resolution, resulting in prohibitive memory demands. To address this, we developed BandHiC, a memory-efficient and user-friendly Python package for organizing and analyzing Hi-C matrices down to sub-kilobase resolution. BandHiC adopts a banded storage strategy that preserves only a configurable diagonal bandwidth of the dense contact matrix, reducing memory usage by up to 99% while maintaining fast random access and intuitive indexing operations. In addition, it provides flexible masking mechanisms to handle missing values, outliers, and unmappable regions, and supports efficient vectorized operations optimized with NumPy, thereby enabling scalable analysis of ultra-high-resolution Hi-C datasets.

## Introduction

High-throughput chromosome conformation capture (Hi-C) [1–4] and its variants, such as Micro-C [5–7], have substantially advanced our understanding of genome architecture by enabling genome-wide mapping of chromatin interactions at progressively higher resolutions. Hi-C data typically measures the interaction frequency between evenly spaced chromatin segments, represented mathematically as a contact matrix, where each bin corresponds to a genomic interval and the bin size defines the resolution of the contact map. Over the past decade, improvements in sequencing throughput and experimental protocols have enabled a dramatic increase in Hi-C data resolution, from early megabase (Mb)-scale maps to kilobase (kb)- and even sub-kilobase (e.g., 500 bp, 250 bp) contact matrices [4,6]. More recently, single-cell Micro-C has achieved contact maps at resolutions as high as 5 kb [8]. These advances in resolution have facilitated the discovery of finer-scale chromatin structures and their regulatory roles in gene expression [3,6,7,9,10].

However, higher-resolution data generated by these assays pose significant computational challenges, particularly in terms of memory consumption [11,12]. For instance, loading a dense Hi-C matrix into Random Access Memory (RAM) at 1 kb resolution for the human genome (∼3 billion base pairs) would require approximately (3 × 10^9^ / 10^3)^)^2^ × 8 bytes = 7.2 × 10^13^ bytes = 72 ter*a* bytes of memory (≈ 66 tebibytes), assuming double-precision floating-point representation (8 bytes per entry). Such memory demands far exceed the capacities of most computational environments. This issue becomes even more pronounced at sub-kilobase or single-cell resolutions, rendering dense matrix representations impractical for many real-world applications. These limitations highlight the need for more memory-efficient data structures tailored for high-resolution Hi-C data.

Many current methods for identifying genome structural patterns, such as chromatin loops and topologically associating domains (TADs), rely heavily on dense matrix representations to facilitate rapid, random data access during computation [13]. Tools, such as TopDom [14], MSTD [15], DeTOKI [16] and SnapHiC [17], exhibit high memory consumption associated with dense matrices, which limits their scalability to higher-resolution Hi-C datasets. Although conventional sparse matrix formats can reduce memory usage, they lack efficient random-access capability, significantly slowing downstream analyses and complicating algorithm design, particularly for tasks requiring frequent element-wise access or submatrix extraction. Other methods, including Mustache [18] and Chromosight [19], attempt to circumvent this limitation by partitioning the full dense matrix into smaller blocks that are loaded into memory on demand. However, these methods introduce added implementation complexity and require careful memory management to maintain computational efficiency.

However, these methods typically leverage high-resolution Hi-C data predominantly within short-range genomic distances (typically within 10 Mb), as chromatin loops and TADs are generally constrained to such scales [3]. Long-range interactions are increasingly sparse at high resolutions and often irrelevant to local structures such as loops and TADs. Thus, focusing on short-range contacts is both computationally and biologically justified. Therefore, there is an urgent need for a novel memory scheme tailored specifically for short-range higher-resolution Hi-C data, one that dramatically reduces memory usage while retaining the fast random-access capabilities of dense matrices.

Built upon NumPy [20,21], a fundamental library for numerical computing in Python, we developed **BandHiC**, a memory-efficient Python package specifically designed for organizing and analyzing short-range contacts of Hi-C data down to sub-kilobase resolutions. BandHiC adopts a banded storage scheme that stores only a configurable diagonal bandwidth of the full Hi-C contact matrix. Importantly, it preserves efficient random-access capabilities by employing a direct index mapping between the banded and dense matrix representations, while supporting familiar NumPy-style indexing semantics (slicing, Boolean array indexing, integer array indexing) to facilitate user-friendly and efficient data access. BandHiC also integrates masking functionality akin to NumPy’s `MaskedArray` module, enabling straightforward handling of gaps, outliers, and other aberrant values in Hi-C matrices. Finally, BandHiC supports diverse numerical operations optimized through NumPy’s efficient vectorized computations, thus offering both memory efficiency and high computational performance essential for practical high-resolution Hi-C data analysis.

## Design and implementation

### Data representation

To address the increasing memory demands posed by high-resolution Hi-C data, we introduce `band_hic_matrix`, the core class implemented in the BandHiC package. Given a Hi-C contact matrix *A* ∈ *R*^*n*×*n*^ at resolution *r*, `band_hic_matrix` retains only the diagonals within a user-defined bandwidth *k*, yielding a compact representation *D* ∈ *R*^*n*×*k*^ (Fig 1). This format ensures that each column in D corresponds to a fixed diagonal of *A*, such that the mapping *A*[*i, j*] = *D*[*i, j* ― *i*] holds for |*i* ― *j*| < *k*.

**Fig 1.**
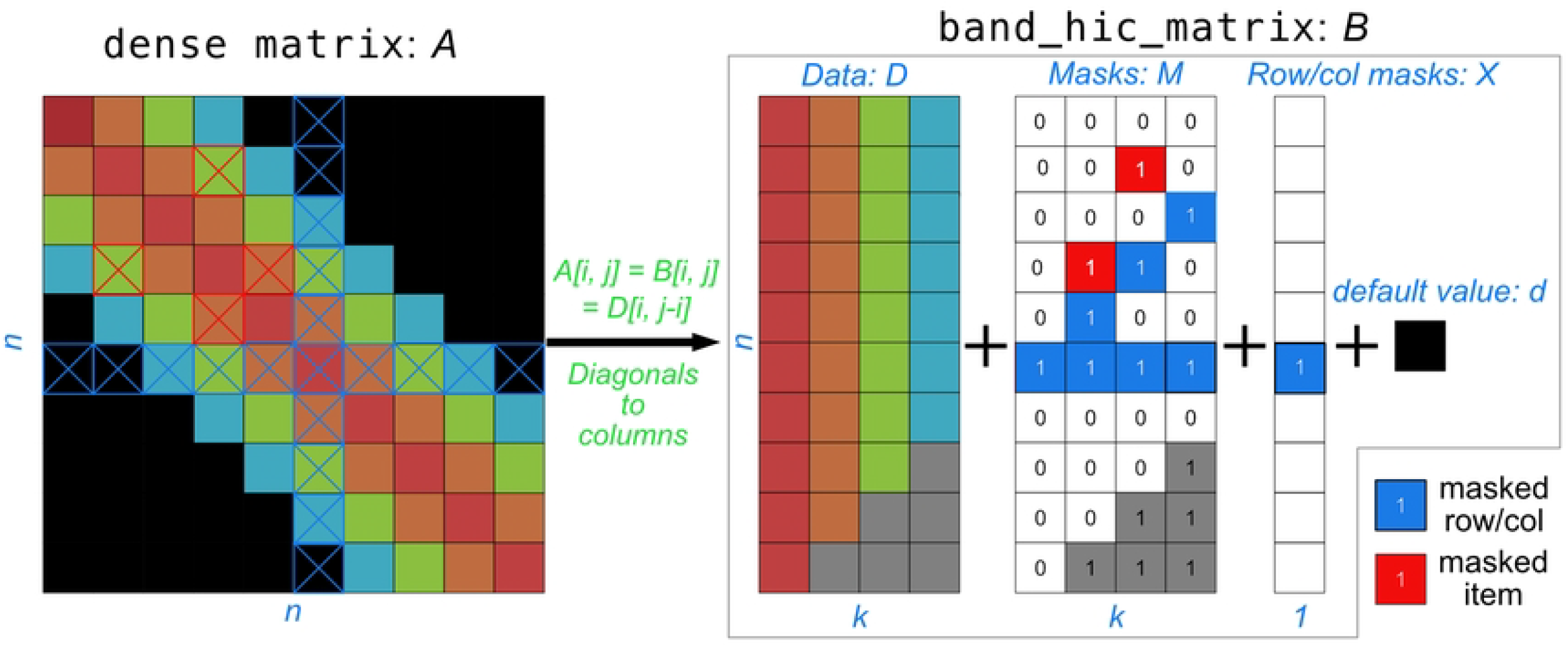
Data model of BandHiC. Schematic illustration of converting a dense symmetric matrix *A* into a banded representation consisting of a data matrix *D*, an element-wise mask matrix *M*, a row/column mask matrix *X*, and a default value *d* for out-of-band entries. Diagonal elements from *A* are reorganized into columns of *D*; *M* marks missing or outlier entries; *X* indicates masked rows or columns.

The memory efficiency achieved by this strategy is substantial. When *k* ≪ *n*, the memory footprint of `band_hic_matrix` is reduced from 𝒪(*n*^2^) to 𝒪(*nk*). For example, assuming a resolution of 1 kb and a bandwidth of 2 Mb (*k* = 2000), the representation of chromosome 1 of the human genome (∼249 Mb) requires 3.7 GiB of memory, less than 1% of the memory required by the dense matrix (∼461.9 GiB). This compression makes high-resolution Hi-C data accessible even on commodity hardware, without compromising the efficiency of random data access.

To further enhance the flexibility of usage, `band_hic_matrix` supports an optional two-layer masking mechanism. An element-wise mask matrix *M* ∈ {0,1}^*n*×*k*^ allows users to selectively ignore missing or outlier contacts, enabling robust statistical estimation on unmasked subsets. Additionally, a bin-level mask *X* ∈ {0,1}^*n*^ supports the exclusion of entire rows or columns, particularly useful for removing repetitive genomic regions lacking valid Hi-C signals. These masking features facilitate downstream tasks such as estimation of average contact intensity at specific genomic distances, while preserving statistical validity.

Lastly, a scalar default value *d* is defined to fill in the undefined entries of *A* not covered by the banded matrix *D*. This default is typically set to 0, consistent with the assumption that long-range interactions are negligibly sparse. The advantage of using default values is that, in addition to treating the `band_hic_matrix` as a full dense matrix for indexing and conversion with a dense matrix, it also allows out-of-band entries to participate in mathematical operations. For example, when adding 1 to the `band_hic_matrix` object, not only are the in-band entries incremented, but the default value representing the out-of-band entries also increases by 1. Together, the components *D, M, X*, and *d* allow for seamless reconstruction of the dense matrix *A* when required. Overall, `band_hic_matrix` provides an efficient, flexible data representation for scalable Hi-C data analysis.

### BandHiC package

BandHiC is distributed as an open-source Python package under the MIT license. It is compatible with Python version 3.11 or higher and can be deployed on Linux and macOS platforms. The BandHiC package relies solely on four dependencies: NumPy, SciPy, cooler, and hic-straw. NumPy and SciPy serve as the fundamental backbone of BandHiC, supporting the construction of the banded matrix data structure and implementing its core computational operations. In addition, BandHiC wraps the file-reading functions of cooler and hic-straw, allowing it to directly read.hic,.cool, or.mcool files as inputs for creating a `band_hic_matrix` object (Fig 2A).

**Fig 2.**
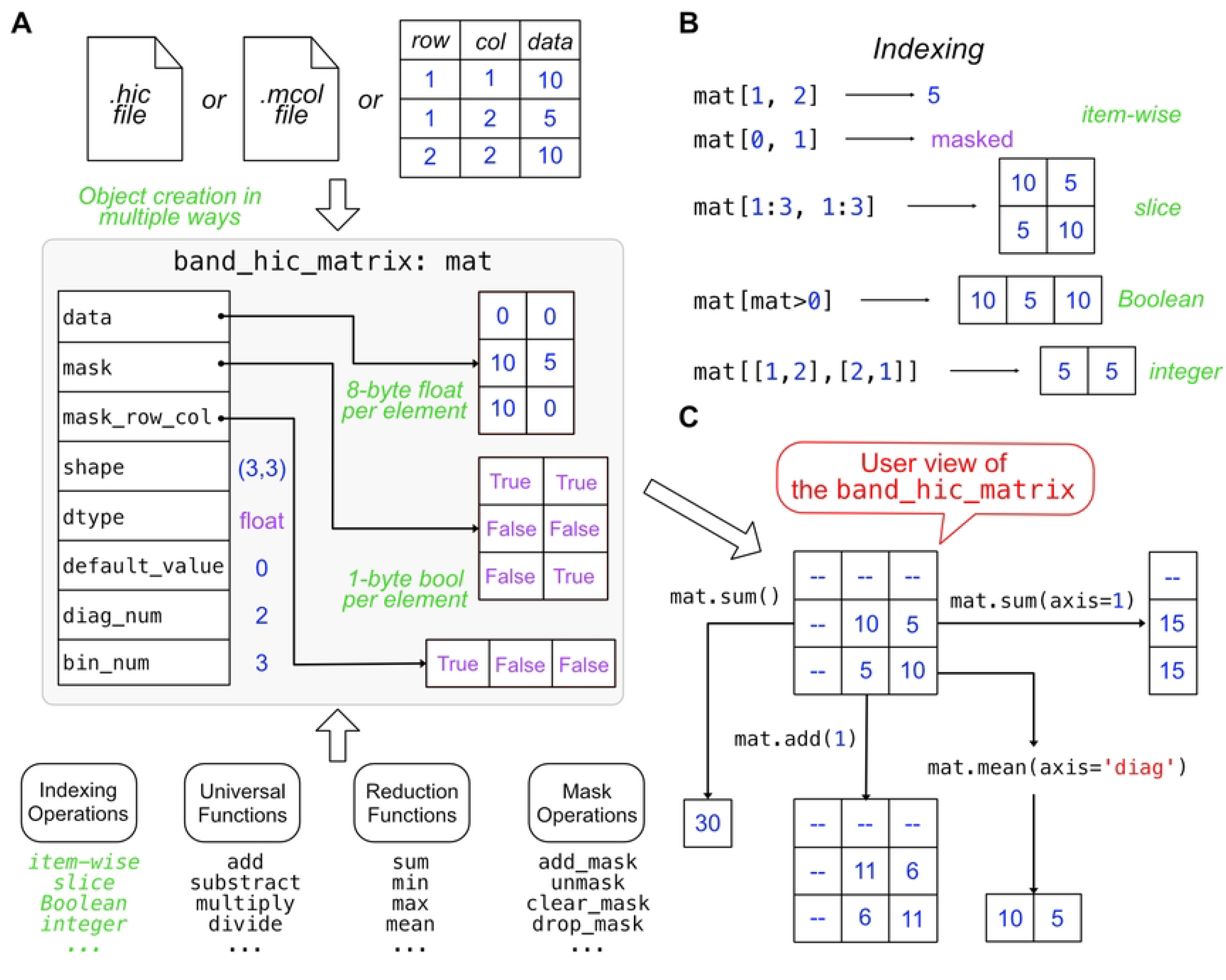
Overview of the BandHiC package. (A) Example of a `band_hic_matrix` object in the BandHiC package. (B) Indexing methods supported by BandHiC. (C) Example of computation methods supported by BandHiC.

BandHiC primarily defines a matrix class, `band_hic_matrix`, which represents Hi-C contact data in a banded matrix representation. Each instance contains a numerical NumPy array of shape `(bin_num, diag_num)`, together with Boolean arrays `mask` and `mask_`row`_col` that record element-wise and row/column-wise exclusions. Elements outside the stored bandwidth are represented by a scalar `default_value`, allowing the matrix to behave as a dense symmetric array while avoiding redundant storage. Objects can be constructed from.hic,.mcool files, or from triplet-form contact records (rows, columns, and contact frequencies) (Fig 2A).

The package fully supports NumPy-style indexing (Fig 2B), including item-wise, slice, Boolean array, and integer array indexing. It also provides a series of methods and functions for constructing, manipulating, and performing computations on the `band_hic_matrix` objects (Fig 2C). The design and implementation of the indexing and computational operations supported by BandHiC are described in detail in the following two subsections.

### Indexing operations

A key feature of `band_hic_matrix` is its direct coordinate mapping between the banded matrix *B* and the full dense matrix *A* (Fig 1). For any pair of genomic loci (*i, j*) satisfying the band constraint: |*i* ― *j*| < *k*, the interaction frequency *A*[*i, j*] can be accessed in constant time via *D*[*i, j* ― *i*]. This mapping ensures random access in 𝒪(1) time, which is critical for performance-sensitive Hi-C analyses, particularly when memory constraints preclude the use of fully dense matrices. 𝒪(1) random access means that any element can be retrieved in constant time, independent of dataset size, because its memory location is computed directly from the index. This property is particularly important for matrix-like data structures such as Hi-C contact matrices, where efficient access to arbitrary entries is essential for large-scale analysis. Data access in `band_hic_matrix` is fully consistent with that of a dense matrix, as each entry is accessed via *B*[*i, j*] = *D*[*i, j* ― *i*] = *A*[*i, j*]. As a result, users can interact with a `band_hic_matrix` object as if it were a standard dense array, without needing to consider the underlying storage details.

In computer science, indexing is the process of locating and accessing elements through their position or key, providing direct and efficient data retrieval compared to sequential search. Leveraging this random-access capability, `band_hic_matrix` supports full NumPy-style indexing semantics, including slicing, Boolean array, and integer array indexing (Fig 2B). Slicing selects contiguous ranges of data, such as *A*[2:5]. Boolean array indexing extracts elements that meet specific conditions, for example *A*[*A* > 0]. Integer array indexing allows arbitrary selection using arrays of integer indices, such as *A*[[0, 2, 4]]. This design allows users to easily query local chromatin contacts and provides a flexible and efficient framework for data manipulation in scientific computing. For instance, a slice operation such as *B*[*i*:*j, i*:*j*] or *B*[*i*:*j*] retrieves a banded submatrix. Combined with the `todense` operation, this enables reconstruction of the dense submatrix for downstream analysis or visualization.

For a `band_hic_matrix` object with a mask matrix, indexing operations return either a `ma.MaskedArray` object or the `masked` constant. In NumPy, a `MaskedArray` stores numerical data together with a Boolean mask that marks missing or invalid entries. This ensures that such values are automatically excluded from computations while maintaining the array structure, making it well-suited for handling incomplete or noisy datasets. In practice, indexing a `band_hic_matrix` behaves as if operating directly on a `MaskedArray`, thereby providing users with considerable flexibility.

Since indexing may refer to elements outside the predefined diagonal bandwidth, such out-of-band entries do not raise errors but are instead filled with a default value (e.g., zero or other user-specified constants). This design further enhances the robustness and flexibility of BandHiC in practical applications.

### Numerical computation

In addition to flexible data access, `band_hic_matrix` also supports a wide range of numerical operations, including element-wise mathematical operations and reduction operations (Fig 2C). BandHiC is built on top of NumPy, which provides the foundation for efficient numerical computation in Python. NumPy offers high-performance, C-optimized universal functions (ufuncs) that perform element-wise operations with support for broadcasting and type casting. By leveraging these ufuncs, BandHiC implements 71 element-wise mathematical operations directly on the `band_hic_matrix` (Table 1). This design eliminates the need for explicit Python loops, ensures full compatibility with the NumPy ecosystem, and significantly improves computational speed. Consequently, BandHiC inherits the scalability and efficiency of NumPy, enabling fast and memory-efficient analysis of large genomic contact matrices. Moreover, NumPy provides interfaces for defining custom array-like objects while maintaining seamless integration with NumPy, enabling BandHiC to implement specialized matrix types efficiently and flexibly. By building on NumPy in this way, BandHiC inherits both its computational efficiency and its flexible programming model, making it well-suited for scalable analysis of large-scale Hi-C data.

**Table 1.** Universal functions that BandHiC supports.

| Function | Description | Function | Description |
| --- | --- | --- | --- |
| absolute | Absolute value | add | Element-wise addition |
| arccos | Inverse cosine | arccosh | Inverse hyperbolic cosine |
| arcsin | Inverse sine | arcsinh | Inverse hyperbolic sine |
| arctan | Inverse tangent | arctan2 | Arctangent of y/x with quadrant |
| arctanh | Inverse hyperbolic tangent | bitwise_and | Element-wise bitwise AND |
| bitwise_or | Element-wise bitwise OR | bitwise_xor | Element-wise bitwise XOR |
| cbrt | Cube root | conj | Complex conjugate |
| conjugate | Alias for conj | cos | Cosine function |
| cosh | Hyperbolic cosine | deg2rad | Degrees to radians |
| degrees | Radians to degrees | divide | Element-wise division |
| divmod | Quotient and remainder | equal | Element-wise equality test |
| exp | Exponential | exp2 | Base-2 exponential |
| expm1 | $\exp(x) - 1$ | fabs | Absolute value (float) |
| float_power | Floating-point power | floor_divide | Integer division (floor) |
| fmod | Modulo operation | gcd | Greatest common divisor |
| greater | Element-wise greater-than test | greater_equal | Greater-than or equal test |
| heaviside | Heaviside step function | hypot | Euclidean norm |
| invert | Bitwise inversion | lcm | Least common multiple |
| left_shift | Bitwise left shift | less | Element-wise less-than test |
| less_equal | Less-than or equal test | log | Natural logarithm |
| log1p | $\log(1 + x)$ | log2 | Base-2 logarithm |
| log10 | Base-10 logarithm | logaddexp | $\log(\exp(x) + \exp(y))$ |
| logaddexp2 | Base-2 version of logaddexp | logical_and | Element-wise logical AND |
| logical_or | Element-wise logical OR | logical_xor | Element-wise logical XOR |
| maximum | Element-wise maximum | minimum | Element-wise minimum |
| mod | Remainder (modulo) | multiply | Element-wise multiplication |
| negative | Element-wise negation | not_equal | Element-wise inequality test |
| positive | Returns input unchanged | power | Raise to power |
| rad2deg | Radians to degrees | radians | Degrees to radians |
| reciprocal | Element-wise reciprocal | remainder | Modulo remainder |
| right_shift | Bitwise right shift | rint | Round to the nearest integer |
| sign | Sign of input | sin | Sine function |
| sinh | Hyperbolic sine | sqrt | Square root |
| square | Square of input | subtract | Element-wise subtraction |
| tan | Tangent function | tanh | Hyperbolic tangent |
| true_divide | Division that returns a float |  |  |

Reduction operations refer to functions that aggregate multiple values into a single result. They can be applied globally to all elements of a matrix, or along specific axes to summarize rows or columns. Examples include `sum`, `min`, `max`, and `mean`. BandHiC supports ten such reduction operations (Table 2), which work along conventional axes (rows or columns) in the same way as NumPy. In addition, BandHiC extends these operations to the diagonal axis—a feature not available in NumPy. This diagonal reduction is particularly useful for Hi-C data, as it allows interaction frequencies to be summarized by genomic distance, thereby supporting distance-dependent normalization and analyses such as distance-decay profiling. All operations remain fully compatible with masked `band_hic_matrix` objects, ensuring robust handling of missing or low-quality data in large-scale Hi-C analysis.

**Table 2.** Reduction functions that BandHiC supports.

| Function | Description |
| --- | --- |
| `sum` | Compute the sum of all elements along the specified axis |
| `prod` | Compute the product of all elements along the specified axis |
| `min` | Return the minimum value along the specified axis |
| `max` | Return the maximum value along the specified axis |
| `mean` | Compute the arithmetic mean along the specified axis |
| `var` | Compute the variance (average squared deviation) |
| `std` | Compute the standard deviation (square root of variance) |
| `ptp` | Compute the range (max - min) of values along the axis |
| `all` | Return True if all elements evaluate to True |
| `any` | Return True if any element evaluates to True |

The implementation of reduction operations in BandHiC is designed to behave equivalently to those on a dense matrix, but without explicitly constructing the dense matrix, which would otherwise consume substantial memory. During computation, BandHiC automatically fills out-of-band entries with the default value, symmetrizes interactions in the lower-triangular part of the matrix, and excludes entries masked by either element-wise or row/column masks.

Taken together, `band_hic_matrix` combines the memory efficiency of a banded storage model with the expressiveness of NumPy’s interface. By mimicking both Numpy’s `ndarray` and `MaskedArray` behaviors, it provides an intuitive and powerful interface for users, substantially lowering the barrier to adoption and enabling seamless integration into existing Hi-C data analysis pipelines. Please refer to BandHiC’s website for a detailed list of all supported functions and tutorials.

## Results

### Usage Examples

BandHiC can serve as an alternative to the NumPy package when managing and manipulating Hi-C matrices, aiming to address the issue of excessive memory usage caused by storing dense matrices with NumPy’s `ndarray`. At the same time, BandHiC supports masking operations like NumPy’s ma.MaskedArray module, with enhancements tailored for Hi-C data. Users can leverage their experience with NumPy when using the BandHiC package, so it is recommended that users have some basic knowledge of NumPy. Here are some code examples providing a quick guide and demonstration of the core functionalities of BandHiC:

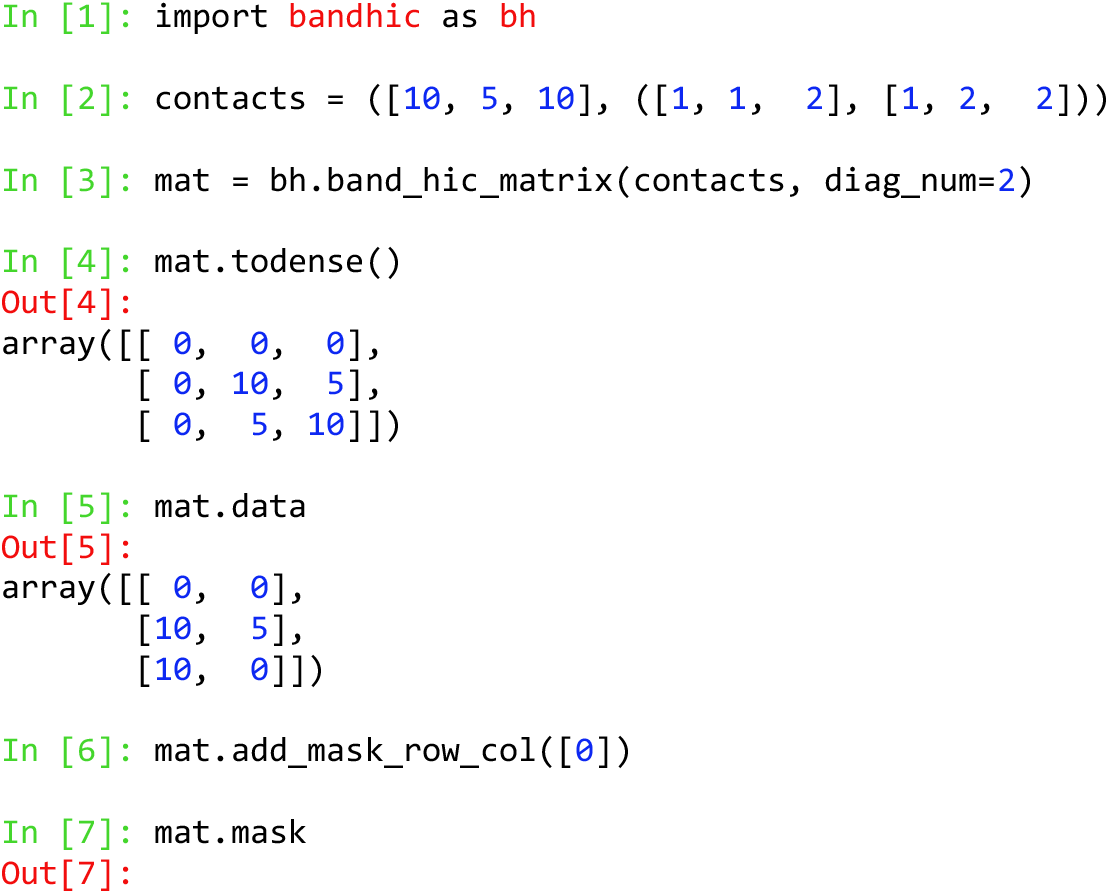
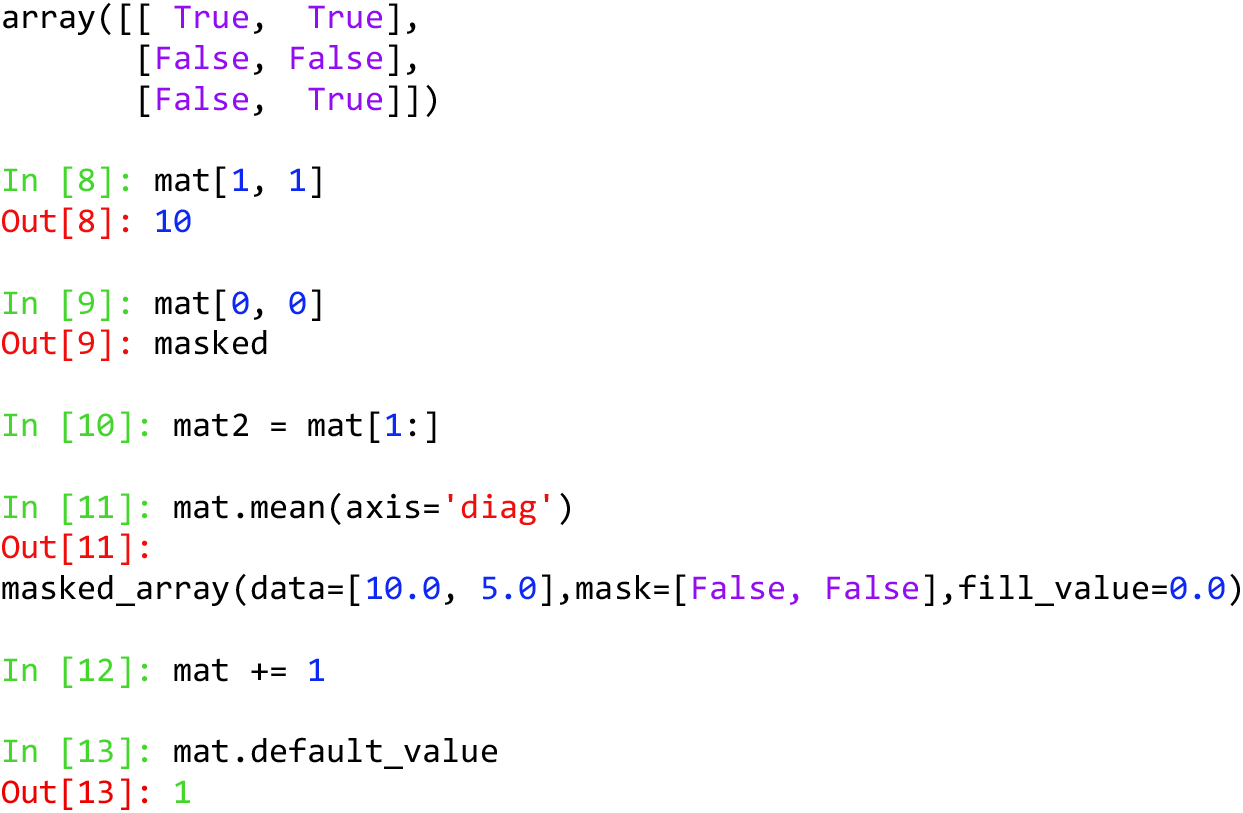

The defined `band_hic_matrix` object `mat` corresponds to the example shown in Fig 2. The example is presented in the IPython-style interactive format, in which “In [n]:” indicates input commands and “Out[n]:” indicates the corresponding outputs.

### Application: Reducing TopDom memory consumption

While the previous section demonstrates the core functionality and syntax of BandHiC, here we evaluate its practical utility by integrating it with the TAD-calling algorithm TopDom [14]. TopDom is an efficient and robust algorithm for detecting topologically associating domains (TADs). It uses a sliding window approach on Hi-C contact matrices to identify local minima in contact intensity, which mark potential TAD boundaries. Owing to its robustness and reproducibility, TopDom was rated as the best-performing TAD detection method in a benchmark study [9]; however, its reliance on dense matrices makes it difficult to apply to sub-kilobase resolution Hi-C data. The original TopDom algorithm was developed in the R language [14]. In BandHiC, we provide a Python implementation of TopDom as a built-in function, ensuring functional consistency with the original method while extending support for both NumPy’s `ndarray` and BandHiC’s `band_hic_matrix` class. This integration not only facilitates direct identification of TADs from high-resolution Hi-C data within the BandHiC but also enables a fair comparison of memory usage and runtime performance between dense and banded representations.

To evaluate the effectiveness of BandHiC, we benchmarked the TopDom algorithm on chromosome 1 of mouse embryonic stem cell (mESC) Micro-C data across multiple resolutions using both dense matrix and `band_hic_matrix`. As shown in Fig 3A, BandHiC substantially reduces memory usage at all tested resolutions. At 1000 bp and 500 bp resolution, the dense matrices require over 72 GiB of memory, exceeding the available system memory (60 GiB RAM and 12 GiB swap space), and causing the program to terminate due to memory allocation failure. In contrast, the `band_hic_matrix` version of TopDom completes successfully, with memory usage of only 5,989 MiB and 23,902 MiB at 1000 bp and 500 bp resolution, respectively. The banded matrix introduces a modest increase in runtime compared to the dense matrix (Fig 3B). This overhead arises not from masking operations (which were disabled for this evaluation) but from index computation performed during element access within the `band_hic_matrix` object. These results indicate that `band_hic_matrix` enables TAD-calling algorithms like TopDom to process high-resolution Hi-C data on a standard personal computer, making large-scale analysis feasible without incurring significant computational cost. Owing to BandHiC’s NumPy-like API, other Hi-C-based pattern identification methods can be reimplemented with minimal modifications to the original code, thereby significantly improving their scalability and adaptability to higher-resolution Hi-C datasets.

**Fig 3.**
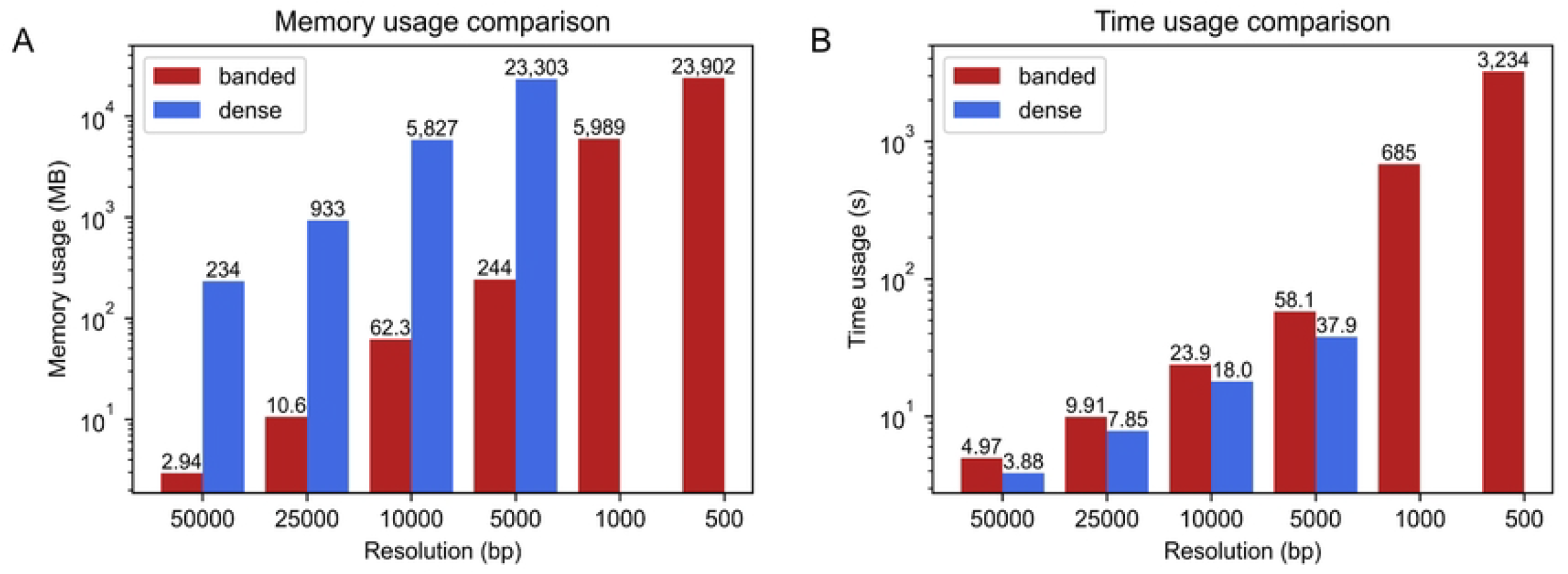
Evaluation of BandHiC. (A) Memory usage comparison between the banded and dense matrices when running TopDom on mouse embryonic stem cell (mESC) Micro-C data (chromosome 1) at various resolutions. (B) Runtime comparison for the same task. At 1,000 bp and 500 bp resolution, the dense representations failed due to memory overflow.

## Availability and Future Directions

The source code of the BandHiC Python package is publicly available on GitHub (https://github.com/xdwwb/BandHiC-Master), and comprehensive documentation is provided at https://xdwwb.github.io/BandHiC-Master/. Installation can be performed conveniently through Python’s pip package manager:

`$ pip install bandhic`

BandHiC alleviates the computational challenges of high-resolution Hi-C analysis through a banded storage scheme that reduces memory usage to ∼1% of dense matrices while preserving constant-time random access. This enables domain-calling algorithms, such as TopDom, to run at sub-kilobase resolution on standard hardware. Seamless integration with the NumPy ecosystem, including support for universal functions, reductions, and Hi-C–specific diagonal operations, facilitates efficient distance-dependent analyses and easy adoption in existing pipelines. Furthermore, BandHiC’s flexible programming model—featuring masking, scalar defaults, and robust handling of noisy or unmappable regions—provides an extensible framework for downstream applications ranging from loop and TAD detection to single-cell Hi-C analysis.

Nevertheless, BandHiC’s current design focuses on short-range, cis interactions, which may limit its applicability for studying long-range or inter-chromosomal contacts that can also carry biological relevance. Future work could address this limitation by introducing hybrid storage strategies that combine banded and sparse formats. In fact, BandHiC can already be applied to single-cell Hi-C analysis by constructing a separate `band_hic_`matrix object for each cell. A more direct and scalable solution would be to extend the underlying `numpy.ndarray` used for banded storage from two to three dimensions, thereby introducing a “cell axis”. This modification would further enhance the utility of BandHiC for single-cell Hi-C data, providing users with a more convenient and powerful tool for large-scale single-cell 3D genome studies. Additionally, integrating BandHiC with visualization tools or downstream analysis pipelines could further broaden its applicability in exploring multi-scale chromatin organization.

Taken together, BandHiC represents both a practical solution to the pressing computational challenges posed by ultra-high-resolution Hi-C data and a flexible foundation for future methodological advances. By combining memory efficiency, computational scalability, and a user-friendly interface, BandHiC has the potential to become a core component of the bioinformatics toolkit for 3D genomics.

## Data availability

Micro-C data for the mouse embryonic stem cell (mESC) line were obtained from the NCBI GEO database (accession number: GSE130275).

## Acknowledgements

We thank all the members of Prof. Gao’s lab for helpful comments.

